# Crosstalk between myosin II and formin functions in the regulation of force generation and actomyosin dynamics in stress fibers

**DOI:** 10.1101/2021.08.15.456175

**Authors:** Yukako Nishimura, Shidong Shi, Qingsen Li, Alexander D. Bershadsky, Virgile Viasnoff

## Abstract

REF52 fibroblasts have a well-developed contractile machinery, the most prominent elements of which are actomyosin stress fibers with highly ordered organization of actin and myosin IIA filaments. The relationship between contractile activity and turnover dynamics of stress fibers is not sufficiently understood. Here, we simultaneously measured the forces exerted by stress fibers (using traction force microscopy or micropillar array sensors) and the dynamics of actin and myosin (using photoconversion-based monitoring of actin incorporation and high-resolution fluorescence microscopy of myosin II light chain). Our data revealed new features of the crosstalk between myosin II-driven contractility and stress fiber dynamics. During normal stress fiber turnover, actin incorporated all along the stress fibers and not only at focal adhesions. Incorporation of actin into stress fibers/focal adhesions, as well as actin and myosin II filaments flow along stress fibers, strongly depends on myosin II activity. Myosin II-dependent generation of traction forces does not depend on incorporation of actin into stress fibers per se, but still requires formin activity. This previously overlooked function of formins in maintenance of the actin cytoskeleton connectivity could be the main mechanism of formin involvement in traction force generation.

**Highlights:**

• Cell traction forces are measured together with visualization of actomyosin flow
• Actin turnover depends on formin and myosin II activities
• Traction forces depend not only on myosin II, but also on formins
• Traction force generation may require formin-dependent cytoskeleton connectivity

## 1. Introduction

Since actin distribution in cultured cells was first visualized by antibody staining [1], the most prominent actin structures identified were bundles of actin filaments known as actin fibers or cables. Further analysis revealed that actin fibers contain numerous actin associated proteins and permitted to classify them into several groups. First dichotomy is between straight actin cables usually called stress fibers [2–5], and curvy or circular actin transverse arcs parallel to the cell edge [6–8]. Stress fibers are always associated with cell adhesions [9–13]. In fibroblast-like cells spread on rigid planar extracellular matrix, these are so-called focal adhesions mediated by integrin family of receptors. The straight stress fibers can be further classified into two groups: the fibers growing from peripheral focal adhesions towards the cell centre known as radial or dorsal stress fibers, and the fibers connecting two focal adhesions known as ventral stress fibers [14, 15]. An important difference between these types of structures is that radial fibers do not contain myosin II [16, 17], while the ventral stress fibers contain myosin II filaments organized into periodically-spaced stacks often considered as primitive sarcomere-like structures [18]. Actin transverse arcs or circumferential actin fibers appear at the cell periphery and also contain myosin II filaments [18, 19]. The actomyosin contraction drives the centripetal movement of these structures [17, 20]. The three types of actin fibers are interrelated, so that radial fibers and arcs are thought to be precursors of the ventral stress fibers. Specifically, a pair of radial fibers associated with focal adhesions are thought to fuse with a part of actomyosin transverse arc giving rise to a ventral stress fibers connecting these focal adhesions [3, 15]. However, in some cases, the stress fibers connecting two focal adhesions can emerge without precursors, via myosin II-driven stretching of the actomyosin network [21].

Myosin II filament-containing actin fibers are the major generators of traction forces exerted by cells on the extracellular matrix. The myosin II-containing transverse arcs transmit forces to focal adhesions through the radial fibers, while the ventral stress fibers generate tension force applied to both focal adhesions associated with them. The actin fibers are embedded into a continuous network of cytoplasmic actin filaments filling the entire cytoplasm. This network contains also myosin II filaments and therefore is contractile [22]. As a result, the traction forces exerted by cells depend not only on contractility of ventral stress fibers and arcs but also on contractility of this intervening actomyosin network.

The important feature of the actomyosin contractile system in non-muscle cells is its high dynamicity. Both actin and myosin II filaments undergo continuous assembly and disassembly with a characteristic turnover time in a range of a minute [18]. In addition, the bulk turnover of the actomyosin system continuously proceeds in the form of centripetal flow of actin structures [6, 18, 19]. This suggests the existence of feedback relationships between the generation of tension forces and the assembly and disassembly processes of the cytoskeletal networks. Such feedback regulation is however insufficiently characterized.

Substantial efforts have been made to understand the mechanical transmission of stress to the substrate at the focal adhesion sites. The general understanding of force generation at the focal adhesion is usually described by “clutch models”, based on dynamic engagement and disengagement of bonds at the focal adhesion site [23–26]. These models, however, do not consider how the mechanical stress is born or dissipated within the stress fibres that are often described for sake of simplicity as load-bearing quasi-static structures. Here, we endeavoured to better characterize the crosstalk between the intrinsic turnover dynamics of molecular components within the stress fibres and mechanical stress propagation.

Our approach was to improve the methods of cell traction force measurement in such a way that simultaneous high-resolution visualization of the dynamics of actomyosin cytoskeleton became possible. Using these correlative measurements, we applied experimental manipulations to disrupt force generation and actin cytoskeleton organization processes. For these manipulations, we focused on pharmacological inhibitors that rapidly affect key components of the actin cytoskeleton rather than genetic knockdown or knockout approaches as the time taken for genetic manipulations to modulate the actin cytoskeleton is longer than the characteristic times for cytoskeletal dynamics we sought to investigate. Careful titration of inhibitor concentrations and treatment time allowed us to avoid any apparent changes in cytoskeleton morphology. Since not all these inhibitors are fully specific, we paid special attention to dissecting the specific effects of interest from the effects on other targets. As a result of these studies, we describe here the basic feedback processes characterizing actin cytoskeleton dynamics.

## 2. Results

### 2.1. Experimental approaches for measuring actomyosin dynamics and cell traction forces

In this section, we summarize the methodological approaches used to assess actomyosin cytoskeleton dynamics and the cell generated traction forces. First, we used photoconvertible β-actin (mEos3.2-β-actin) from Addgene/Michael Davidson lab for assessment of actin incorporation in the ventral stress fibres and focal adhesions [27]. In this set up, we illuminated a circular region (6.5 μm diameter) in the central part of the cell expressing mEos3.2-actin for three seconds to induce green to red photoconversion of labelled actin. Both F-actin and G-actin were photoconverted. Red actin diffused across the cytoplasm to be gradually incorporated into the structures of interest over 5-20 minutes (Fig.1 A-C, Supplementary Figure1, Movie 1). This approach allowed us to investigate the effects of a variety of inhibitors as well as application of stretching forces on actin incorporation. We also used the same construct to label the micron-sized spots on the stress fibres using illumination of cells along parallel lines perpendicular to the stress fibre array. Such ‘zebra’ labelling permitted us to follow the flow of actin filaments along the stress fibres (Fig. 1D, Movie 2). Additionally, we analysed the movement of myosin II filaments along the stress fibres using super resolution Structured Illumination Microscopy (SIM) of cells expressing GFP-fused myosin II regulatory light chain (GFP-MLC). As shown previously, this type of microscopy makes it possible to visualize individual myosin filaments [18]. To quantitively assess the parameters of the actin and myosin II filament flow, we used kymograph analysis for myosin II filaments.

**Figure 1:**
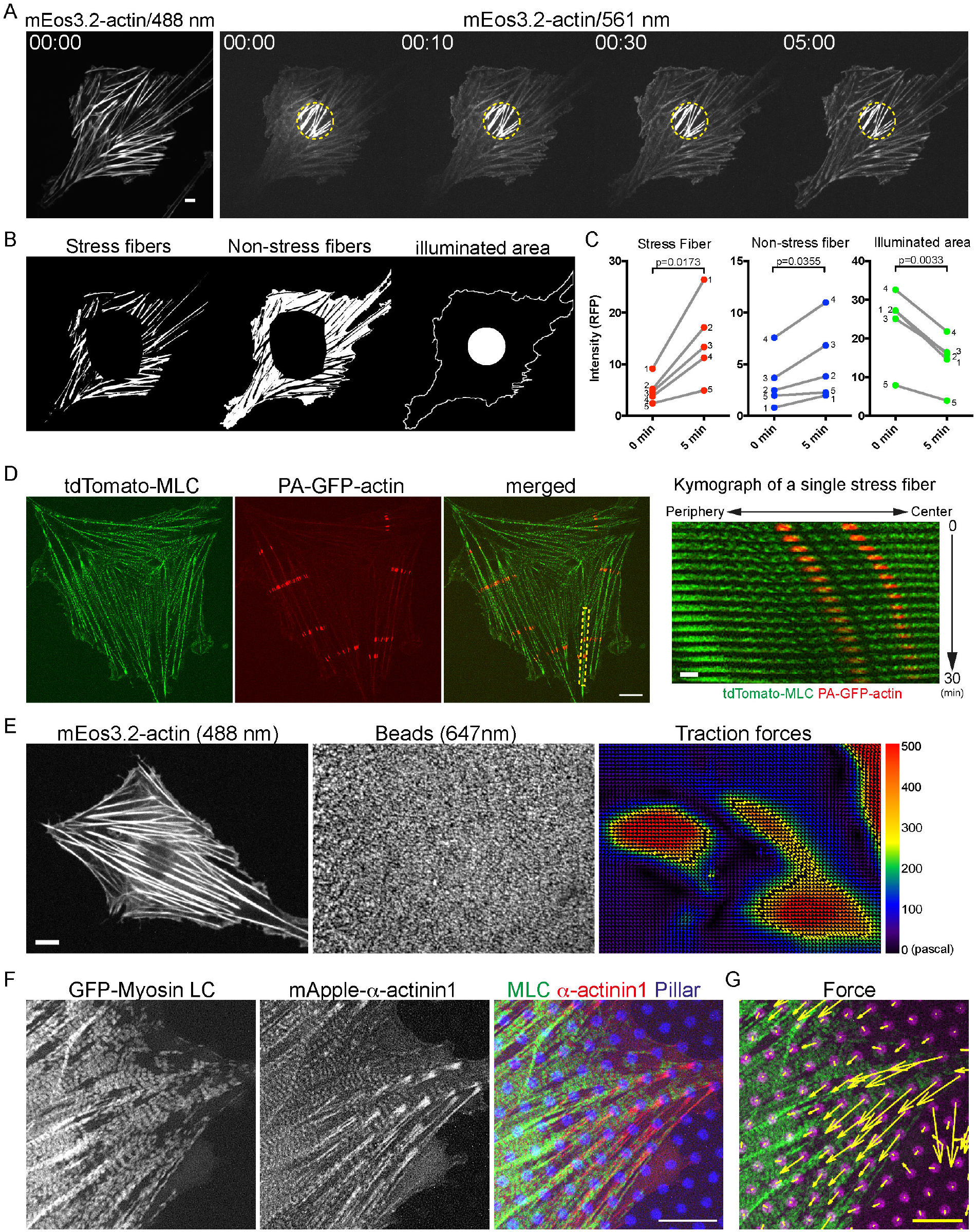
Experimental approaches to assessment of actin dynamics and traction forces. (A) The REF52 cell transfected with photoconvertible actin (mEos3.2-β-actin) is shown in GFP channel (488 nm) before photoconversion (left) and in RFP channel (561 nm) at different moments after photoconversion of the circle in the center (marked in yellow) of the cell by illumination at 405 nm for 3 seconds. Note the gradual incorporation of photoconverted actin into the stress fibers. (B) Masks used for the quantification of the fluorescence intensity of stress fibers (left) or in the space between the stress fibers (right) were obtained by binarization of GFP channel image. The central part of the cell surrounding the illuminated area is removed. (C) Fluorescence intensity of photoconverted actin (RFP channel) immediately after (0 min) and 5 min following photoconversion in the stress fiber region (red), the non-stress fiber area (blue), and central illuminated area (green) of the five cells (numbered). The p-values for the significance of the differences between 0 and 5 minutes time points are calculated by paired two-tailed student *t-test*. The complete curves showing fluorescence dynamics in these five cells are shown in Supplementary Figure 1. (D) SIM images of REF52 cells show myosin filaments (tdTomato-MLC) and illuminated spots of actin filaments (PA-GFP-actin). Kymograph in the right panel shows the actin and myosin flow in the stress fiber in the yellow dotted rectangle (merged image in the left panels). Bar in the right panels, 2µm. (E) Measurement of traction forces of REF52 cells by the traction force microscopy. A representative REF52 cell expressing mEos3.2-β-actin (488nm, the left panel) plated on silicon membrane with fluorescent beads (647nm, the middle panel). Tension magnitude map of the same cell is shown in the right panel (Traction forces). (F) Myosin II filaments, α-actinin1 and traction forces in REF52 cell on micropillar array. SIM images of myosin light chain (left, GFP-MLC) and α-actinin1 (middle, mApple-α-actinin1); merged image of MLC in green, α-actinin1 in red, and fluorescent pillars in blue is shown in the right. (G) The traction force vectors in the same cell calculated from pillar displacement are shown as yellow arrows. Scale bars in A, B, D (right), E and F, 10µm. Scale bar for the force vectors in G, 30 nN.

To combine assessment of actin and myosin II filament flow with measurements of traction forces cells exert on substrates, we followed the approach introduced in the previous study [28] with significant technical improvements. Two methods were used to measure traction force. First, we modified traction force microscopy to use very thin (7 μm) PDMS layers coated with fibronectin and decorated with attached 40 nm fluorescent fiducial beads as a substrate for cells [29]. We used the algorithm described in Tseng et al [30] for computation of traction forces based on bead displacement measurements (Fig. 1E). These thin PDMS layers enabled the simultaneous use of super resolution SIM to visualize myosin II filaments in these cells at 120 nm resolution. Second, we directly measured traction forces using optical distortion-free micropillar arrays made of an elastomer (My-134 polymer, Mypolymer Ltd.) with a refractive index matching that of the growth medium [31]. This method of the traction force measurement was also compatible with SIM microscopy. Actin and myosin dynamics could thus be assessed together with forces exerted by individual stress fibers (Fig. 1F, Movie 3).

### 2.2. The turnover of actin in stress fibers is regulated by tension forces generated by myosin II filaments

We used incorporation of photoconvertible actin to characterize actin turnover in ventral stress fibers. We measured the kinetics of incorporation of mEos3.2-β-actin, photoconverted by illumination of circular area in the cell centre, into the fraction of ventral stress fibres located outside the photoconverted area (see Fig. 1A). In initial experiments, we analyzed separately the mEos3.2-β-actin incorporation dynamics into 3μm length terminal segments corresponding to the focal adhesions and the rest of the stress fibres (Fig. 2A). We found that photoconverted mEos3.2-β-actin rapidly incorporated into focal adhesions and stress fibres alike, albeit with different kinetics (Fig. 2B-D). Actin incorporated homogenously along the entire fiber and did not show prominent advection from the focal adhesion sites. We conclude that both ventral stress fibres and associated focal adhesions undergo rapid turnover and continuously incorporate new actin. In our subsequent analysis, we measured the integral incorporation of the photoconverted mEos3.2-β-actin along the entire length of the stress fibers including focal adhesions.

**Figure 2:**
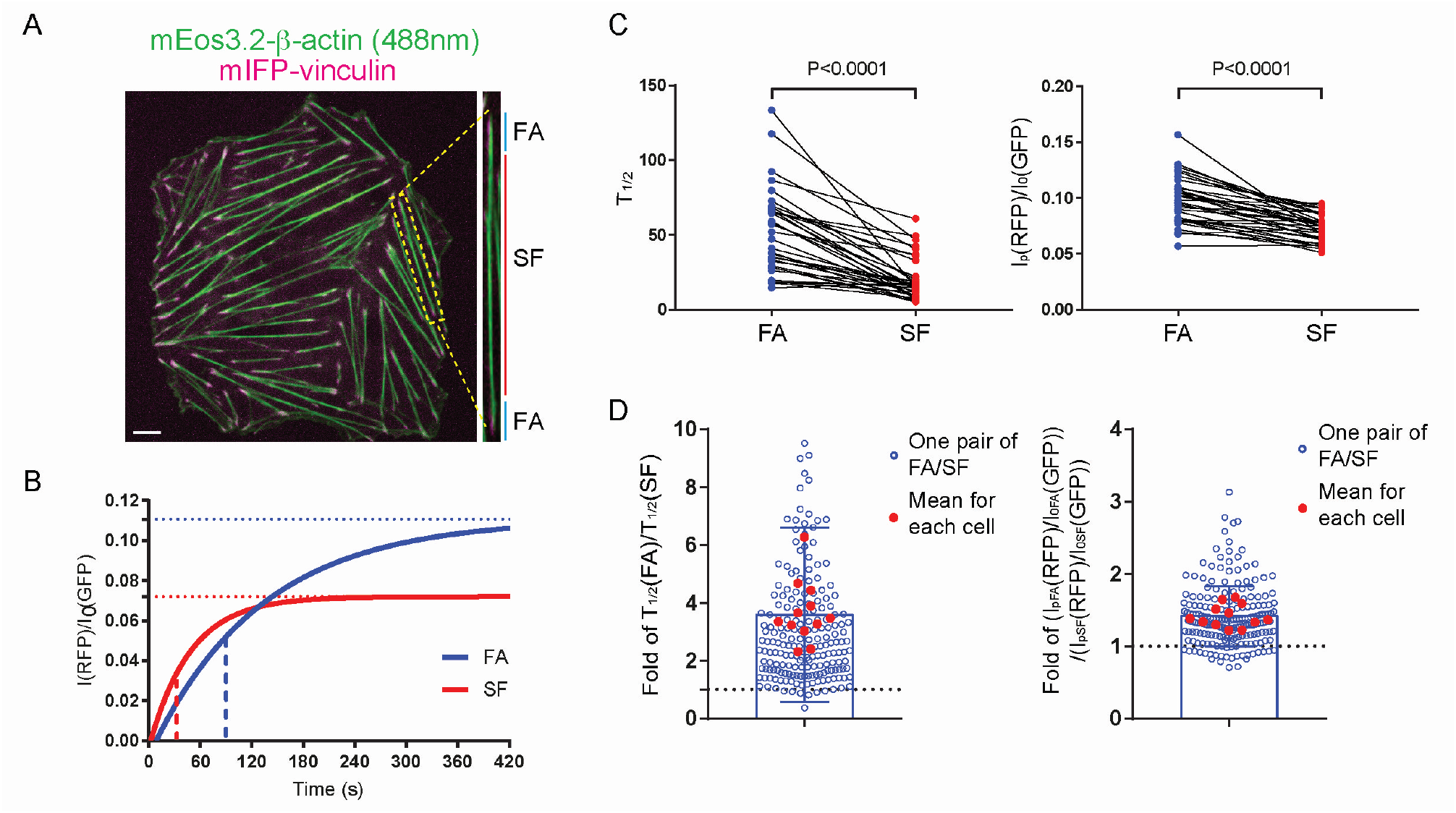
Incorporation of photoconverted actin into focal adhesions and stress fibers. (A) A representative REF52 cell shows stress fibers (non-converted mEos3.2-β-actin, green) and focal adhesions (mIFP-vinculin, magenta). Scale bar, 10 µm. Enlarged image of the stress fiber in the yellow rectangular box is shown in the right. 3 µm length focal adhesion region (FA) and stress fiber proper region (SF) are indicated. (B) Schematic graph showing typical normalized curves of photoconverted actin incorporation I(RFP)/I_0_(GFP) where I(RFP) is fluorescence intensity at RFP channel (photoconverted) and I_0_(GFP) is fluorescence intensity at the initial moment at GFP channel (non-photoconverted). Incorporation into focal adhesion (blue curve) and stress fiber (red curve) regions are characterized by different maximal (plateau) levels (horizontal dotted lines) and half times (marked by vertical broken lines showing half-maximal fluorescence). (C, D) Comparisons between photoconverted actin incorporation into focal adhesion and stress fibers proper regions of the same stress fiber at 10 minutes after photoconversion. (C) Half time (T_1/2_ , left) and plateau levels (I_p_(RFP)/I_0_(GFP), right) are shown for all stress fibers and associated focal adhesions in single cell. Lines connect the paired data points. n = 34 pairs. p-values calculated using paired two-tailed Student’s *t*-test. (D) The ratio values between half times (left) and plateaus (right) characterizing actin incorporation into paired focal adhesion region (FA) and stress fiber region (SF). Blue circles represent the ratios for individual SF/FA pairs, while red dots represent the mean ratios for individual cells. n = 202 pairs, N = 12 cells. Bars indicate mean ± SD for individual SF/FA pairs. Note that majority of the values are more than 1 (dotted lines) showing that both parameters are higher for focal adhesions than for stress fibers proper.

We further assessed the effects of a series of inhibitors on incorporation of photoconverted actin into ventral stress fibers. The doses of the inhibitors used did not produce any apparent changes in the organization of actin (and myosin) during 10-20 minutes of treatment as assessed by spinning disk confocal microscopy with 60X oil immersion objective (see Fig. 3A, 488nm).

**Figure 3:**
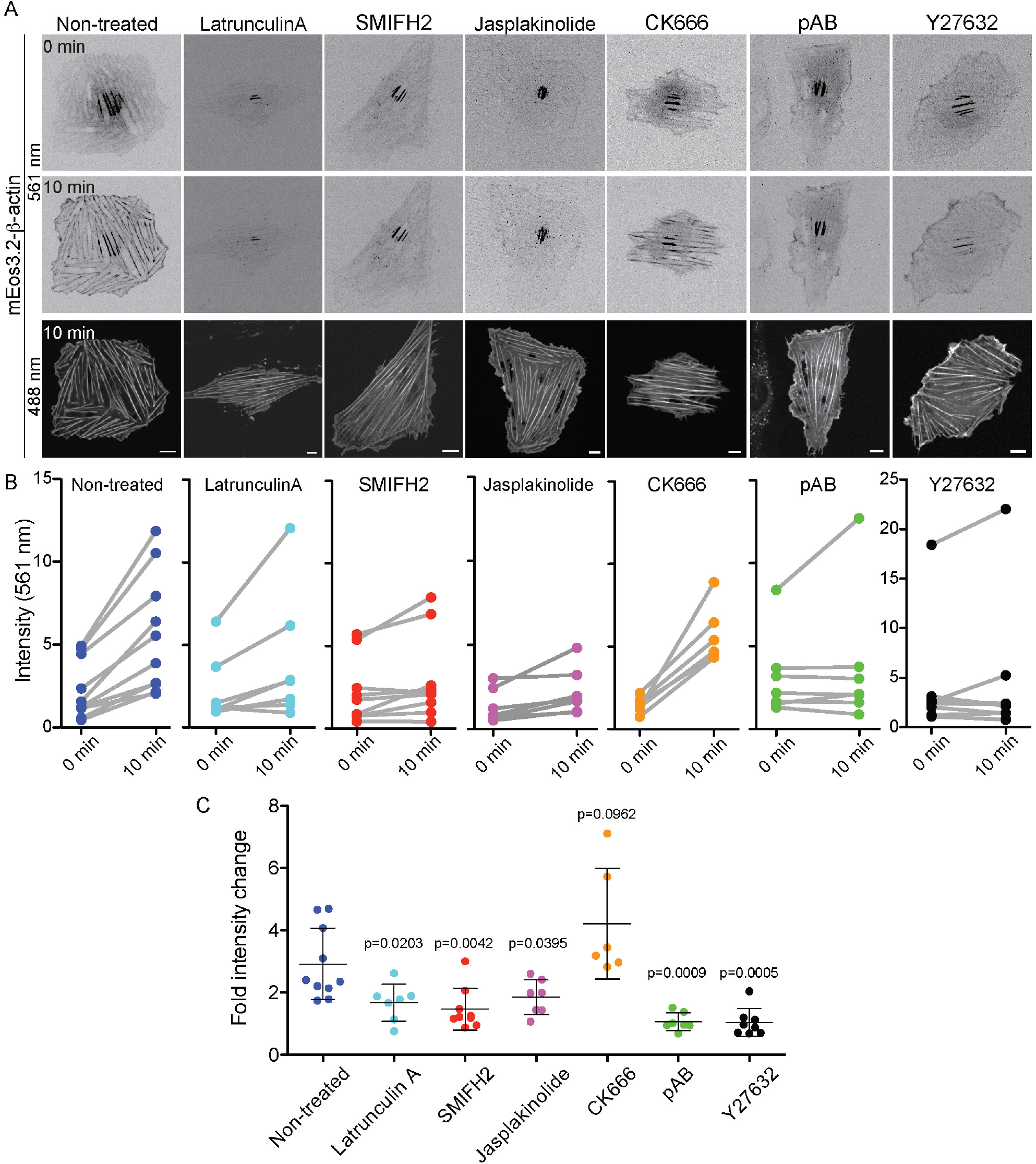
The incorporation of photoconverted actin into stress fibers/focal adhesions depends not only on actin polymerization but also on myosin II activity. (A) Images illustrating incorporation of photoconverted actin in the actin-containing structures of cells pretreated with 30µM SMIFH2 for 5min, 100nM latrunculin A for 30min, 100 nM jasplakinolide for 30 min, 100nM CK666 for 30min, 80µM pAB for 5min or 20µM Y27632 for 5min. Upper and middle rows show inverted images of photo-converted mEos3.2-actin immediately after (upper row) and 10 min following (middle row) photoconversion as visualized by illumination with 561nm light, respectively. Lower row shows the entire actin structures of the cells visualized by non-photoconverted mEos3.2-actin upon illumination with 488nm light at the 10 min time point. Note strong actin incorporation into stress fibers in non-treated and CK666-treated cells but not cells treated with other inhibitors. Scale bars, 10µm. (B) Graphs of the photoconverted actin incorporation (fluorescence intensity at 561 nm illumination) into stress fibers (FA and SF regions together) immediately after (0 min) and 10 min following photoconversion of mEos3.2-β-actin in non-treated cells and cells treated with the drugs, as indicated. Each dot represents incorporation into all stress fibers in one single cell segmented by mask as in Figure 1B, C. (c) Fold changes in the fluorescence intensity of photoconverted mEos3.2-β-actin (ratio between intensity at 10 min and 0 min after photoconversion) in non-treated and inhibitor-treated cells. Each dot represents the ratio values for individual cells. The p values characterizing the differences between mean values for non-treated and drug-treated cells were calculated by an unpaired two-tailed student *t-test*.

Latrunculin A, which inhibits actin polymerization by sequestering monomeric actin and accelerating filament depolymerization [32, 33], completely suppressed incorporation of photoconverted mEos3.2-actin into the stress fibers (Fig. 3). The same effect was observed upon treatment with 100nM jasplakinolide, which stabilizes actin filaments preventing their turnover [34, 35] and therefore also reduces the pool of monomeric actin. Complete suppression of photoconverted actin incorporation was observed also upon treatment with 30μM of SMIFH2, (Fig. 3) which is known to interfere with formin family protein interaction with actin [36] as well as with activity of several types of myosins [27] . However, treatment with a specific inhibitor of Arp2/3 dependent actin polymerization, CK666 [37, 38] did not affect incorporation of the photoconverted mEos3.2-β-actin into the stress fibers even at 100 μM concentration (Fig. 3).

Remarkably, inhibition of myosin II ATPase activity by non-phototoxic derivative of blebbistatin, para-amino blebbistatin (pAb)[39], also completely blocked the incorporation of mEos3.2-actin into the stress fibers, before any changes in the integrity and density of the stress fibers were detected. In addition, we used an inhibitor of Rho kinase (ROCK), Y27632, which is known to suppress myosin light chain phosphorylation and trigger the depolymerization of myosin II filaments [18, 40, 41]. 10 minutes of Y27632 treatment preserved the overall structures of stress fibers, but the incorporation of mEos3.2-actin into the stress fibers was fully blocked (Fig. 3). Thus, inhibition of myosin II ATPase activity and myosin light chain phosphorylation interferes with incorporation of photoconverted β-actin into focal adhesions and stress fibers.

To investigate whether external mechanical forces affect actin incorporation into focal adhesions and stress fibers, we plated cells on stretchable PDMS substrate coated with fibronectin and examined the effect of periodic substrate stretching on the incorporation rate. We designed a custom-made device referred to as a cell stretching dish (Fig. 4A), European Patent application n. PCT/EP2018/053477 (Publication Date 23.08.2018). The substrate underwent biaxial 10 % stretching at 1 Hz frequency for 10 minutes (Fig. 4A, B) on the inverted microscope stage. Comparison of fluorescence intensity of photoconverted mEos3.2-β-actin in cells imaged 10 and 20 minutes after photoconversion revealed that periodic stretching enhanced incorporation of photoconverted actin into stress fibers (Fig. 4C, D). Pre-treatment with SMIFH2, which inhibited incorporation of photoconverted actin under static condition (Fig. 3A, B), did not prevent actin incorporation into stress fibers of periodically stretched cells (Fig. 4E, F). Remarkably, pre-treatment with myosin II inhibitors (para-amino blebbistatin, or Y27632) before periodic stretching resulted in drastic changes to the actin cytoskeleton (Fig. 4E), specifically a strong decrease in fluorescence intensity and number of stress fibers, and formation of lamellipodia. The remaining stress fibers did not incorporate photoconverted actin (Fig. 4E, F).

**Figure 4:**
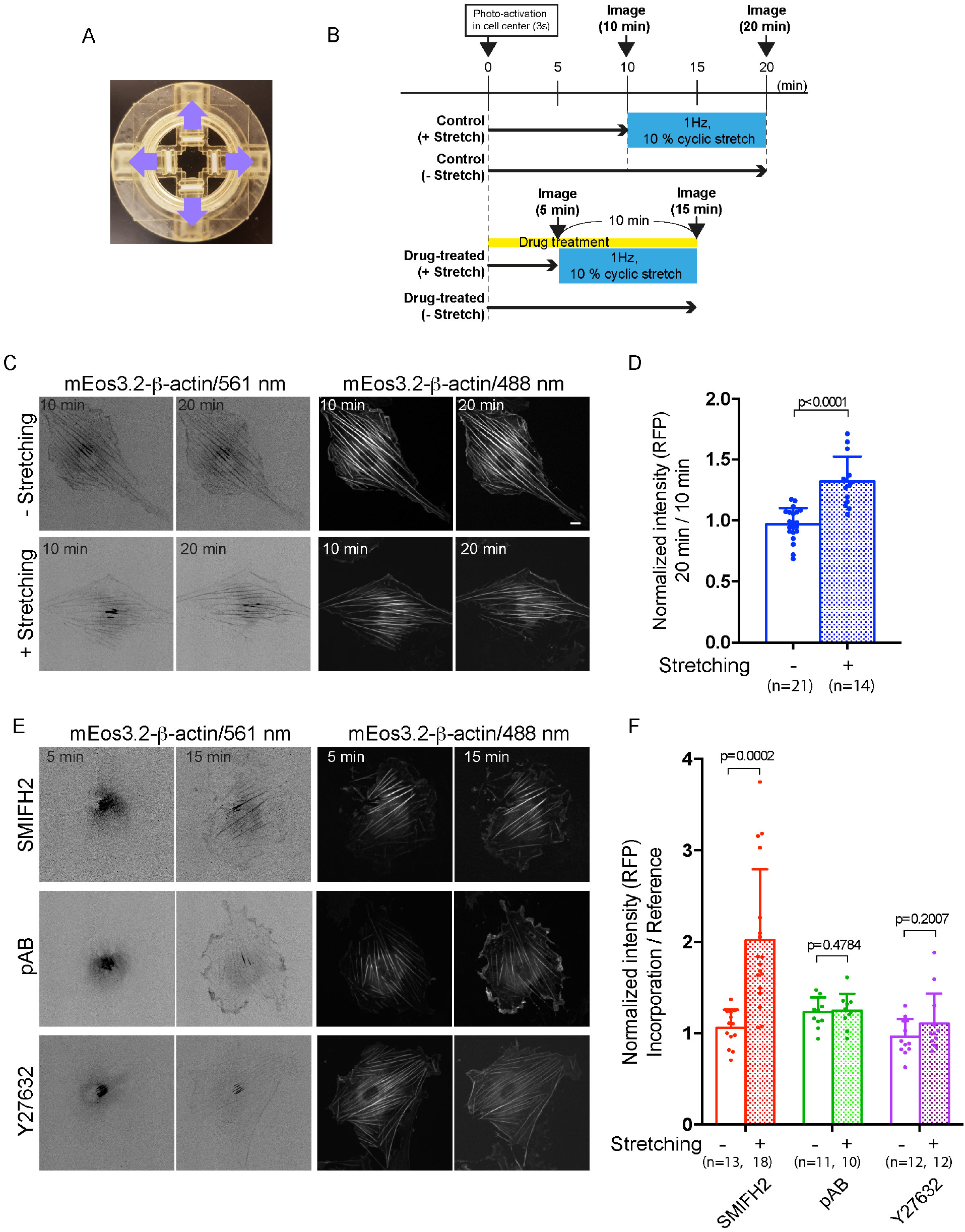
The external forces affect actin incorporation into stress fibres. (A) A top view image of the stretching device. A silicon membrane for cell stretching is located in the center of the device (dark area). Four arms, which are simultaneously driven by an electric motor move centrifugal (purple arrows) and back to apply biaxial cyclic stretching (1 Hz, 10% for 10 min). A schematic diagram shows timetable of stretching and actin incorporation assay under different conditions. In experiment without drug treatment (two upper time arrows), the cells were illuminated at their center and incubated for 10 min (when actin incorporation reached the plateau level), followed by another 10 min with (upper arrow) or without (lower arrow) cyclic stretching. The images were taken at 10 and 20 min following photo-activation, in the beginning and the end of the stretching period. In drug-treated groups (two lower time arrows), the drugs were added simultaneously with the start of photo-activation. The stretching period started at 5 min and ended at 15 min following the photoconversion. The images were taken at the beginning and the end of the stretching period. (C) Fluorescence microscopy images of the photoconverted (561 nm) and non-converted (488 nm) mEos3.2-β-actin in REF52 cells in the stretching device. Photoconverted actin images are inverted. The actin incorporation is increased in cells after cyclic stretching as compared to that in cells before stretching, whereas no change was observed in cells incubated without stretching between 10min and 20 min following photoconversion. (D) The ratio between fluorescence intensity of photo-converted actin in cells after and before stretching. Dots represent the ratio values for individual stretched or non-stretched cells. p-value calculated using two-tailed Student’s *t*-test is indicated. Error bars indicate ± SD. (E) Representative images of the photoconverted (561 nm) and non-converted (488 nm) mEos3.2-β-actin in cells treated with 30µM SMIFH2, 75µM pAB or 20µM Y27632 before and after cyclic stretching. Cyclic stretching rescued the actin incorporation into stress fibers/focal adhesions in cells treated with formin inhibitor SMIFH2, but not in cells treated with myosin II inhibitors pAB and Y27632. (F) The ratio of fluorescence intensity of photo-converted actin in drug treated cells before and after stretching period. Dots represent the ratio values for individual cells. n, number of cells measured under each condition. p-values calculated using two-tailed Student’s *t*-test are indicated. Error bars indicate ± SD. Scale bars, 10µm.

### 2.3. Involvement of formins in the regulation of cell traction forces

To reveal the possible feedback between actin turnover and myosin II-dependent force generation, we then checked how inhibitors of actin polymerization affect cell traction force generation. We used latrunculin A, jasplakinolide, CK666, and SMIFH2 compounds characterized above. As mentioned previously, in addition to formin inhibition, SMIFH2 can inhibit myosin family proteins including non-muscle myosin IIA [27]. In this study, we carefully compared the effects of SMIFH2 and myosin II inhibitors (pAB and Y27632) on traction force generation to dissect out the two activities of this drug and reveal the role of formins in the regulation of cell contractility.

Traction force microscopy performed 10 min after adding 0.1 μM latrunculin A did not reveal any decrease in cumulative traction force per cell area (Fig. 5G, Supplementary Figure2). However, detailed measurements of traction forces generated by individual stress fibres using pillar micro arrays revealed that prolonged latrunculin A treatment (20 minutes) slightly decreased the average value of stress fiber-generated traction forces, but the effect was still not prominent. Detailed inspection of latrunculin A treated cells revealed local defects in actomyosin cytoskeleton organization (oval ‘holes’, Supplementary Figure 3A, C, Movie 5). Measurements of local pillar deflection dynamics revealed that latrunculin A treatment reduced the traction forces only in regions with these local defects of actomyosin organization (Supplementary Figure 3B and C). The overall inhibition of actin polymerization upon treatment with latrunculin A was detected earlier than appearance of these defects (Fig. 3A, B) and by itself did not reduce the traction forces. Treatment of cells with actin filament stabilizing drug jasplakinolide, which strongly reduced actin incorporation into stress fibers as shown above (Fig. 3), did not reduce the traction forces (Fig. 5G, Supplementary Figure2B). Similarly, treatment of cells with Arp2/3 inhibitor CK666 (100μM of for 10 min) did not induce any noticeable changes in traction forces (Fig. 5G).

**Figure 5:**
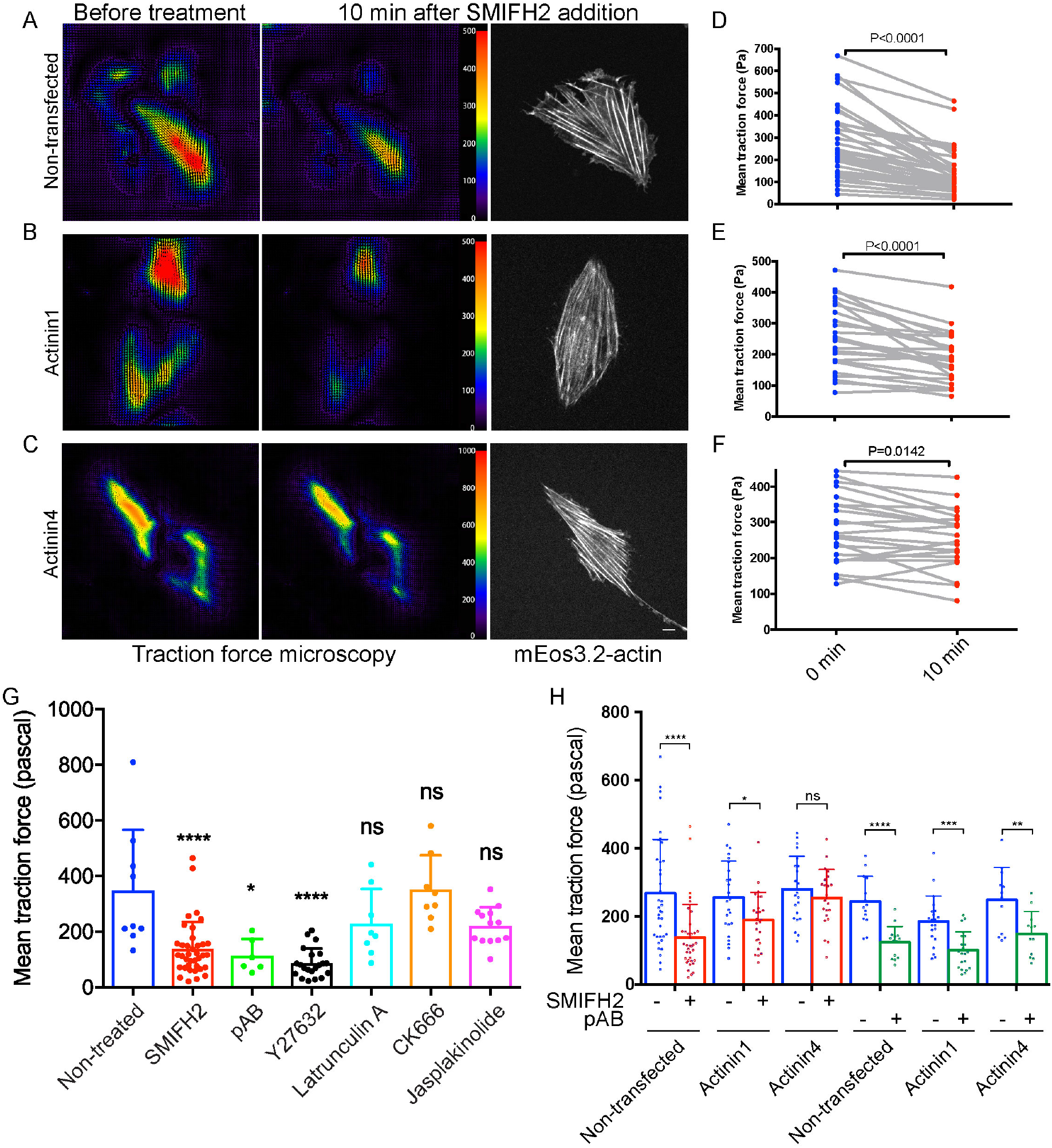
Effects of SMIFH2 and other inhibitors on traction force generation by control and α-actinin1 and -4 overexpressing cells. (A-C) Heat maps of traction forces in cells on elastic substrate at 0 min and 10 min following the addition of 30 µM SMIFH2. (A) - control cell, (B) - cell overexpressing α-actinin1, (C) - cell overexpressing α-actinin4. The overall organization of actin stress fibers (non-photoconverted mEos3.2-β-actin) does not change at 10 min following the drug treatment. Scale bar, 10µm. Measurements of the mean traction force drop in individual cells treated as indicated is shown in (D), (E), and (F), respectively. The mean traction force values in the same cell at 0 min and 10 min following SMIFH2 addition are connected with grey lines. The p-values characterizing the effect of SMIFH2 treatment in control and α-actinin1 and -4 overexpressing cells were calculated using paired two-tailed student *t-test*. Note that drop of traction forces induced by SMIFH2 treatment was significantly less prominent in α-actinin4 overexpressing cells than in control ones. The overexpression of α-actinin1 was less efficient if at all. (G) The mean magnitude of traction forces generated by control REF52 cells and cells treated with indicated drugs (30 µM SMIFH2 for 15min, 100 nM latrunculin A for 40 min, 100 nM jasplakinolide for 40 min, 100 nM CK666 for 40 min, 75 µM pAB for 15 min, and 20 µM Y27632 for 15min). Note that only SMIFH2, pAB and Y27632 reduced the traction forces while other inhibitors do not. (H) The mean magnitude of traction forces in non-transfected and α-actinin1 and -4 transfected REF52 cells treated with SMIFH2 or myosin inhibitor pAB for 10 minutes. Reduction of traction forces by SMIFH2 was prevented by overexpression of α-actinin4 (and in a much lesser degree by α-actinin 1), while the inhibitory effect of pAB was not prevented by either α-actinin 4 or 1 overexpression. The dots in G and H graphs show the magnitude of traction forces in individual cells. Error bars indicate ± SD. The p-values calculated using an unpaired two-tailed student t-test are shown by asterisk representation: * p < 0.05, ** p < 0.01, *** p < 0.001, **** p < 0.0001. ns: p > 0.05.

Unlike latrunculin A, jasplakinolide and CK666, treatment with SMIFH2 resulted in a drop in traction forces generation. This inhibition was observed both in experiments where the forces were measured using traction force microscopy (Fig. 5G), or measured by pillar deflection (Supplementary Figure 2B). Of note, the concentration of SMIFH2 used in these experiments (30 μM) was lower than the concentration of this drug that inhibits myosin II A [27].

In addition to myosin IIA activity, cell traction force is regulated by the degree of actin filament cross-linking mediated, in particular, by α-actinin family proteins [42, 43]. Here, we examined how increasing α-actinin1/4 levels affected contractility inhibition by SMIFH2 and blebbistatin (pAB) (Fig. 5A-F, H). In agreement with a previous study [43], we found that overexpression of α-actinin4 increased the fraction of non-polarized cells lacking ventral stress fibers. However, the percentage of elongated cells exhibiting a system of parallel ventral stress fibers remained around 50 %. In these elongated cells, overexpression of α-actinin4 by itself did not significantly affect the mean cell traction force (Fig. 5H). However, α-actinin4 overexpression remarkably prevented the traction force decrease induced by SMIFH2 treatment (Fig. 5C, F, H), but not the traction force decrease induced by blebbistatin (Fig. 5H). Overexpression of α-actinin1 seems to act in a similar way albeit its effect was much weaker (Fig. 5B, E, H). These data demonstrate that inhibition of traction forces by SMIFH2 can be prevented by increase of actin filament cross-linking by α-actinin. The difference between SMIFH2 and myosin II inhibition strongly suggests that the loss of tension cannot be attributed to an off-target effect of SMIFH2 on myosin II activity [27], and hence that formins are directly involved in the maintenance of traction forces.

### 2.4. Comparison of effects of SMIFH2 and blebbistatin (pAB) on actin and myosin II flow

To further elucidate the dynamics of actin and myosin II in the stress fibers, we compared the effects of blebbistatin and SMIFH2 on the flow of actin and myosin filaments in the segments of stress fibers adjacent to the focal adhesions. The actin was labeled by photoactivation of PA-GFP-β-actin; the average distance between photoactivated actin spots and the ends of stress fibers (focal adhesion) was 5-10 μm (Fig. 6A, Movie 6). The myosin II filaments were labeled by td-Tomato-MLC and the flow was estimated by the movement of fluorescent spots corresponding to myosin filament clusters. Observation of the flow in control cells revealed that both actin spots and myosin filaments moved centripetally along stress fibers with an average velocity of 0.13 μm/min (Fig. 6B), in the fibers which did not retract from the focal adhesion sites. Addition of para-amino blebbistatin (pAb) 5 minutes before starting flow observation revealed a drop in the average velocity of both actin and myosin filaments to 0.05 μm/min (Fig. 6B). In some cases, flow stopped entirely by 15 min following the start of the observation. Unlike blebbistatin, addition of SMIFH2 5 min before starting observation induced a pronounced increase in the velocity of centripetal movement of both actin and myosin filaments. The average flow velocity increased to 0.6 μm/min and remained at this level for observation period of 15 min (Fig. 6B). Simultaneous treatment with SMIFH2 and pAB stopped the flow similar to treatment with pAB alone (Fig. 6E).

**Figure 6:**
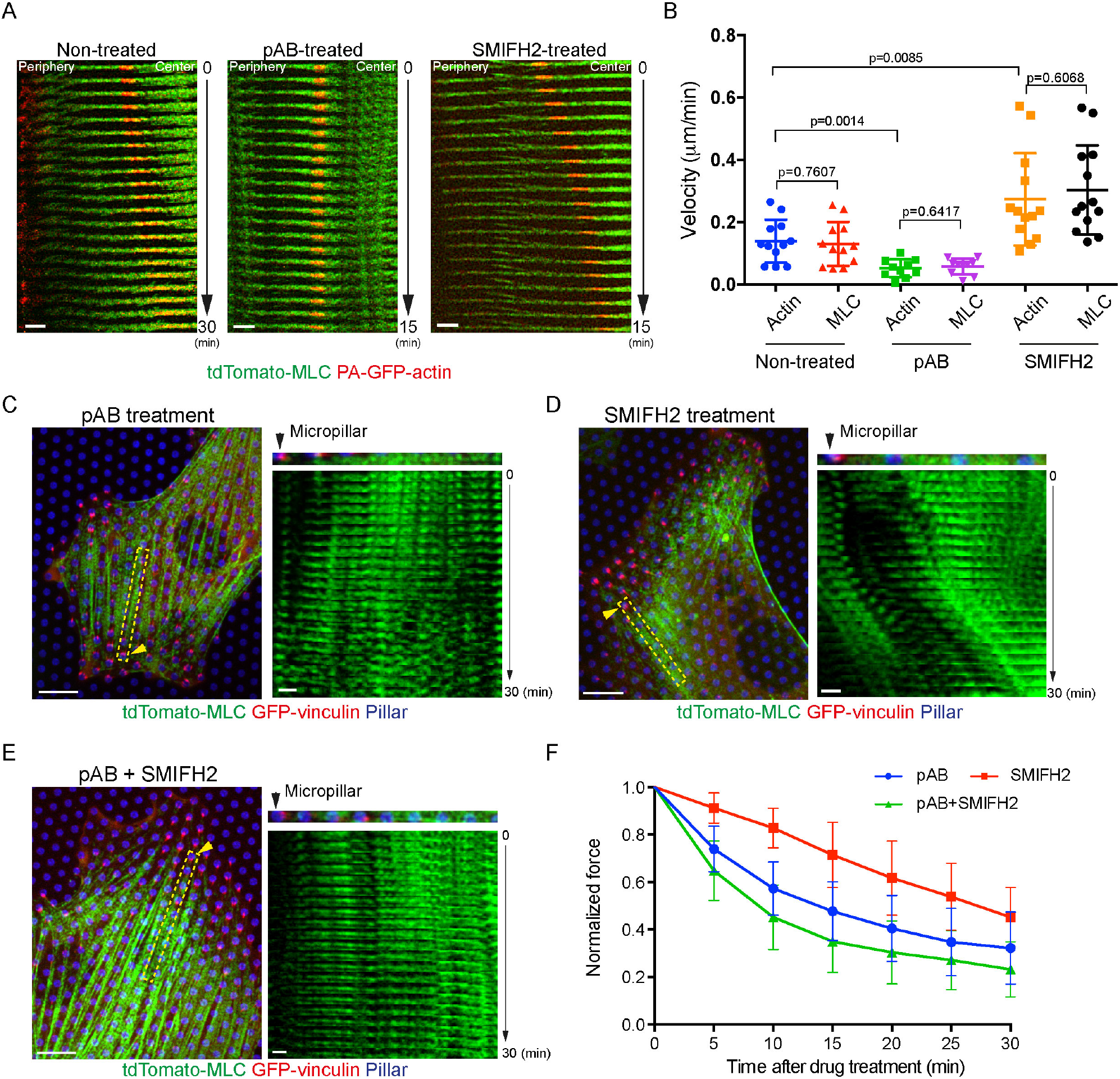
The effects of SMIFH2 and blebbistatin (pAB) on actin and myosin II flow and force generation in stress fibers. (A) Kymographs showing the flow of myosin filaments (green, tdTomato-MLC) and actin filaments (red, photoconverted PA-GFP-β-actin) in individual stress fibers in non-treated cell (left), and cells treated with 100 μM pAB (middle) and 25 μM SMIFH2 (right). Focal adhesions are located on the left side of each kymographs (marked as ‘Periphery’). Bars, 2 μm. Duration of the kymographs for untreated cell – 30 min, for treated cells – 15 min. (B) Graph showing the velocities of movement (calculated from displacement per 15 min) of photoconverted actin spots and adjacent myosin II filaments along stress fibers. The p-values calculated using an unpaired two-tailed student t-test are indicated. Both actin and myosin II filaments moved with similar speed. pAB treatment inhibited the flow, whereas SMIFH2 accelerated velocity of both actin and myosin II filaments in the stress fibers. (C-E) Distribution of myosin II filaments (green, tdTomato-MLC) and focal adhesions (red, GFP-vinculin) and kymographs showing the flow of myosin II filaments in REF52 cells treated with 100 μM pAB (C), 25 μM SMIFH2 (D) or the mixture of 100 μM pAB and 25 μM SMIFH2 (E) in cells on micropillar arrays (the tips of pillars are shown in blue, Atto647N-fibronectin). In each panels, left image shows the entire cell, while kymograph on the right shows the movement of myosin II filaments in the stress fiber marked by yellow dotted box in the left image. Arrows show the position of the micropillars, associated with the boxed stress fibers via focal adhesions (red). Bars in the left images - 10 μm, in the right images - 2 μm. Similarly to (A) and (B), the treatment with pAB stopped the myosin II filament flow, while the treatment with SMIFH2 accelerated it. Combined treatment with pAB and SMIFH2 stopped the flow. (F) The measurement of the traction forces upon treatment cells with the drugs. The traction forces were calculated by deflections of single micropillars associated with individual stress fibers in cells treated with 25µM SMIFH2 (red line, n=13 stress fibers, 4 cells), 100 µM pAB (blue line, n=30 stress fibers, 5 cells) or the mixture of 25µM SMIFH2 and 100 µM pAB (green line, n=28 stress fibers, 4 cells). The values were normalized by the traction forces at the initial time point. Error bars indicate ± SD. Note the difference in decay rate between SMIFH2 treated cells and the cells treated with blebbistatin (pAB) or the mixture of SMIFH2 and pAB.

Comparison of the changes in myosin flow and traction forces induced by pAB and SMIFH2 in single stress fibers (Fig. 6C-F) revealed a similar inhibition of traction forces 30 min following addition of either 25 µM SMIFH2 or 100 µM pAB. Kymographs of myosin movement at approximately 10-15 micrometers from the end of a stress fiber associated with a pillar demonstrated the absence of myosin flow in the presence of pAB (Fig. 6C, Movie 7) and increase of myosin flow upon treatment with SMIFH2 (Fig. 6D, Movie 8), similarly to the data shown in Fig.6A. The combined treatment with pAB and SMIFH2 led to cessation of the flow as in the case of treatment with pAB alone (Fig. 6E, Movie 9). Thus, either stopping or accelerating myosin flow can accompany a decrease in traction force.

## 3. Discussion

The aim of this study was to elucidate the relationship between the traction forces generated by ventral stress fibers and the actomyosin turnover inside these structures. We found that actin incorporation into stress fibers depends on the forces generated by myosin II. At the same time, the traction forces exerted by the stress fibers depends not only on myosin II, but also on formin activities.

We applied several methods to visualize and quantify the dynamics of actomyosin cytoskeleton in cultured fibroblasts. To quantify actin turnover we used a photoconvertable β-actin construct which allowed us to follow actin incorporation into stress fibers and focal adhesion structures. In addition, we used photoconvertible actin to measure bulk flow of actin along the stress fibers. In our experiments, we measured the movement of micron-sized patches of photo-converted actin (mEos3.2- or PA-GFP-) along stress fibers. These observations permitted us to study the actin flow in the stress fibers, compare it with myosin filament flow (measured directly by SIM imaging of individual myosin filaments), and investigate the effects of actin and myosin II inhibitors on this process.

The limitation of this approach is a lack of information concerning functional activity of the photoconvertable fusion actin construct. It is known that expression of GFP-actin can affect actin dynamics *in vivo* [44] and interfere with some cell functions [45]. Moreover, labelled yeast actin which could incorporate into Arp2/3 dependent actin patches did not incorporate into formin dependent contractile rings [46, 47]. Interestingly, human neutrophil-like HL-60 cells demonstrated normal migration after complete substitution of endogenous β-actin with a GFP fusion construct of β-actin [48].

We found that photoconverted mEos3.2-β-actin rapidly incorporated not only into lamellipodia and focal adhesions, but also along the entire length of the stress fibers. Detailed comparison of mEos3.2-β-actin incorporation into focal adhesions and stress fibers revealed some kinetic differences. The incorporation into the stress fibers was characterized by shorter half time (T _1/2_) but lower plateau level (I_p_) as compared to incorporation into focal adhesions. These kinetic differences may reflect the differences in composition of nucleating and elongating proteins in these structures. For example, focal adhesions are enriched in actin filament-elongating VASP/Ena proteins [49–52] and can contain vinculin-associated components of Arp2/3 actin nucleating complex [53]. Moreover, different formin family proteins could differentially localize to the focal adhesions and stress fibers.

Using this methodology, we studied how different inhibitors of actomyosin force generation affected incorporation of photoconvertable actin into focal adhesions/stress fibers. It is well known that focal adhesion assembly is a mechanosensitive process which can be activated by application of myosin II generated forces or by exogenous mechanical forces [54, 55]. A surprising finding is that the actin incorporation into entire stress fibers can also be efficiently suppressed by inhibition of myosin regulatory light chain phosphorylation (via suppression of Rho kinase by Y27632), as well as by inhibition of myosin II ATPase (by blebbistatin photo-insensitive derivative pAB). Both these inhibitors significantly reduced the level of photoconvertable actin incorporation into stress fibers and focal adhesions. These findings are consistent with our observation that external forces applied to cells through periodic substrate stretching activated actin incorporation into focal adhesions and stress fibers.

Since focal adhesion plaques consist of actin filament bundles [12, 13], the enhanced actin incorporation observed in our experiments could be one of the mechanisms underlying the process of force-dependent focal adhesion assembly. Our finding that incorporation of photoconvertible actin into stress fibers depends on myosin II-driven and external forces reveals a new facet in understanding stress fiber turnover and demonstrates the existence of a complex mechano-regulation feedback for this process. In particular, actin dynamics within the stress fiber does not originate solely from incorporation into focal adhesion sites but occurs over the entire length of the stress fibers.

Incorporation of actin into stress fibers is regulated at different levels [56]. One class of proteins that could be responsible for the force dependence of actin incorporation into focal adhesions and stress fibers are the formin family proteins, which are known to be major nucleators and elongators of actin filaments [57, 58]. Previous studies have clearly demonstrated that mechanical forces stimulate formin mDia1-dependent actin polymerization *in vitro* [59–62]. There are also evidences of mechanosensitivity of actin polymerization by other formins [63–65]. At this stage, it is difficult to elucidate whether mechanosensitivity of actin incorporation into focal adhesions and stress fibers is mediated by formins. Knockdowns of individual formins often result in disorganization of the entire system of focal adhesions and stress fibers and even distort myosin filament organization [18]. In addition, the redundancy between formins is another complication for using a knockdown approach. The only available method to rapidly inactivate the majority of formins in the cell is to use a small molecular weight inhibitor of formin FH2 domain (SMIFH2) introduced as a specific pan-formin inhibitor [36]. Indeed, in our experiments SMIFH2 completely blocked the incorporation of photoconvertable actin into stress fibers and focal adhesions at a concentration lower than that required for its off-target effect on myosin II activity [27].

Our experiments showed that effects of SMIFH2 and myosin filament inhibitors (Y27632 and para-aminoblebbistatin) on actin incorporation are not identical. While application of external stretching forces did not prevent the inhibitory effect of Y27632 and blebbistatin on incorporation of photoconverted actin into focal adhesions and stress fibers, it partially rescued the effect of SMIFH2. This may suggest that function of formins is dispensable for the force-dependent activation of actin incorporation and further emphasizes the role of myosin II in this process. It is possible that myosin II not only generates contractile forces or participates in transmission of external forces but enhances actin filament turnover due to its severing function [66–69]. This effect could in principle be potentiated by application of an external stretching force. Another non-alternative possibility could be that myosin II filaments generate pulling forces recruiting actin filaments from the surrounding intervening actin network [22, 70, 71] into the stress fibers. Further studies are needed to elucidate these mechanisms.

We used two independent methods to measure cell contractile forces. First, we used an improved traction force microscopy method based on measurements of displacement of micron sized beads attached to a thin elastic PDMS film covered with fibronectin [29]. Second, we used the arrays of elastomeric pillars for cell spreading and calculated traction forces by measurements of pillar deflections. The elastomer used for fabrication of the pillars had a refractive index similar to that of the growth medium which improved the optical quality of the images [31]. Thus, our measurements of cell traction forces were compatible with simultaneous SIM super-resolution imaging of actomyosin structures in the cells.

The measurement of traction forces exerted by cells on the substrate permitted us to dissect the roles of myosin II filaments and formins in this process. As predicted, inhibition of myosin light chain phosphorylation or ATPase activity led to significant decrease of traction forces. The treatment with either Y27632 or pAB for 15 min led to a two-fold drop in traction forces. However, treatment with latrunculin A at the concentration which efficiently prevented actin incorporation into the stress fibers did not significantly reduce traction forces. The reduction of actin incorporation by jasplakinolide treatment also did not reduce the traction forces. At the same time, SMIFH2 treatment reproducibly and efficiently decreased traction forces. A previous study showed that the concentration of SMIFH2 required for inhibition of myosin IIA *in vitro* is relatively high [27] and exceeded the concentration used in our traction force experiments. Moreover, we detected an important difference between effects of SMIFH2 and blebbistatin/Y27632 on the traction forces. Upon overexpression of actin filament cross-linking protein α-actinin4, the effect of SMIFH2 was essentially abolished while effects of blebbistatin/Y27632 were not changed. This suggests that SMIFH2-induced decrease in traction forces is not due to inhibition of myosin IIA activity. This SMIFH2 effect on traction forces is not also due to inhibition of actin polymerization, since neither latrunculin A nor jasplakinolide, which reduce incorporation of actin into stress fibers, inhibit traction forces.

To explain SMIFH2 effect on traction forces, we considered our previous work which showed that this drug can efficiently detach formins from the actin filament [72]. Thus, it can be suggested that the mechanism of SMIFH2-induced suppression of traction forces is related to a rapid disconnection of actin filaments from the formin molecules. Formin molecules can multimerize [73, 74] and can be part of molecular complexes (asters, nodes, vertices) connecting the plus ends of actin filaments together [75–78]. In particular, formins play an important role in the structural organization of muscle sarcomeres [79–81]. In addition, some formins demonstrate significant actin cross linking function [82, 83], which in principle could be inhibited by SMIFH2. Altogether, treatment with SMIFH2 suppresses the cell traction forces neither by its effect on myosin II, nor by its effect on actin polymerization, but most probably due to reduction of the actin cytoskeleton connectivity. This idea is consistent with our observation that overexpression of the α-actinin4 crosslinker, which can restore actomyosin network connectivity, partially abolished the SMIFH2 effect. We hypothesize that reduction of actin cytoskeleton connectivity interferes with transmission of forces generated by myosin filaments along the stress fibers.

Our studies of actomyosin flow dynamics also revealed the difference between the effects of SMIFH2 and the myosin II inhibitor pAB. While inhibition of myosin II activity, as expected, stopped the flow of actin and myosin II filaments along the stress fibers, addition of SMIFH2 surprisingly accelerated this flow. This accelerated flow can however be stopped by addition of myosin II inhibitor pAB. We hypothesize that disruption of actin network connectivity in the stress fibers by SMIFH2 induced the rapid myosin II-dependent movement but blocked force transmission to focal adhesions resulting in inhibition of traction forces. Thus, reduction of traction forces can be associated with either interruption of actin flow along the fibers (as in the presence of myosin inhibitors), or with increase of the flow (as in the presence of SMIFH2).

In summary, this study revealed the relationship between myosin II-driven force generation, actin filament polymerization, and actin network connectivity in the dynamics of actomyosin flow and generation of traction forces in stress fibers. We have shown that myosin II activity is required for actin incorporation along the stress fibers. Formins are needed not only for actin incorporation but also for the actin network connectivity, which is necessary for transmission of myosin II-generated forces to focal adhesions.

## 4. Materials and Methods

### 4.1. Cell culture, transfection and drug treatment

The immortalized rat embryo fibroblasts (REF52 cells) cell line [84] were cultured in Dulbecco’s modified Eagle’s medium (DMEM; Invitrogen, 11965092) supplemented with 10% heat-inactivated fetal bovine serum (FBS; Invitrogen, 10082147) and 1% penicillin/streptomycin (Invitrogen, 15070063) at 37℃ and 5% CO_2_. The cell line were regularly tested for mycoplasma contamination by MycoAlert PLUS Mycoplasma Detection Kit (Lonza, LT07-703). Cells were transiently transfected with expression vectors using jetPRIME transfection reagent (Polyplus transfection, 114-15) or electroporation (Neon transfection system, Life Technologies) in accordance with the manufacturer’s protocols. The expression vectors used were: mEos3.2-β-Actin, PA-GFP-β-actin, mApple-α-actinin1, α-actinin4-mCherry, tdTomato-MLC (myosin regulatory light chain), mIFP(monomeric infrared fluorescent protein)-vinculin, GFP-vinculin (Michael W. Davidson group collection, The Florida State University, Tallahassee, FL, USA, kindly provided by Dr. P. Kanchanawong, MBI), α-actinin1-mCherry (gift from C. Otey, University of North Carolina, Chapel Hill, NC, USA), and GFP-MLC [85] (a gift from W. A. Wolf and R. L. Chisholm, Northwestern University, Chicago, IL, USA).

Pharmacological treatments were performed using the following concentrations of inhibitors: 25-30 μM for SMIFH2 (Sigma-Aldrich, S4826), 100nM Latrunculin A (Sigma-Aldrich, L5163), 100nM Jasplakinolide (Sigma-Aldrich, J4580), 100nM CK666 (Sigma-Aldrich, SML0006), 10-20μM for Y-27632 dihydrochloride (Sigma-Aldrich, Y0503), 75-100µM Para-aminoblebbistatin (pAB; Optopharma, DR-Am-89).

### 4.2. Live cell imaging

W1-spinning-disc confocal unit (Yokogawa-Gataca systems) mounted on Nikon Eclipse Ti-E inverted microscope (Nikon) with Perfect Focus System 3 and iLas 2 system controlled by MetaMorph software (Molecular device), supplemented with the objective Plan Apo 100x oil NA1.45 or 60x 1.20 NA CFI Plan Apo Lambda water immersion (Nikon) and scientific complementary metal–oxide– semiconductor (sCMOS) camera Prime95B (Photometrics) was used in all experiments. Temperature and CO_2_ level were maintained at 37 °C and 5%, respectively using LCI CU-501 Temperature controller and LCI FC-5N CO_2_ mixer (Live Cell Instrument, Republic of Korea).

Super-resolution SIM imaging was performed using the W1 confocal unit coupled with the live super-resolution (Live-SR) module (spinning disk based structured illumination super resolution [86], Gataca Systems). Laser lines wavelength 405, 488, 561 and 647nm were used.

One hour prior to imaging, L-15 medium without phenol-red (Leibovitz, Sigma-Aldrich) with 10% FBS was added to the transfected REF52 cells.

### 4.3. Actin incorporation assay

REF52 cells transfected with a vector encoding photoconvertible mEos3.2-β-actin were plated overnight on fibronectin-coated 35 mm glass bottom dishes (Iwaki, 3930-035) or on fibronectin coated silicon membrane in stretching device (see below). A circle region with r = 6.5µm in the center of the cell (avoiding the nucleus) was illuminated for 3 seconds by 150 cycles of illumination with 0.06mW 405nm laser for photoactivation. After the illumination, the cells were imaged at 1 min interval using GFP- and RFP-channels visualizing the fluorescence of non-photoconverted and photo-converted mEos3.2-β-actin, respectively.

For the quantitative analysis of actin incorporation, three different masks, namely stress fiber region, cytoplasmic region and lamellipodia region were made using non-photoconverted actin image (GFP-channel) to measure intensity changes in specific regions of cell by ImageJ. The central circular region where the illumination was performed was excluded from the masks and the intensity changes in this region were measured separately.

The intensity of GFP and RFP channels over time were measured. For each type of masks, the intensity of the fluorescence at each given moment was normalized per initial intensity.

For the experiments comparing actin incorporation processes in focal adhesion and stress fiber regions, the images were stabilized by Image Stabilizer (ImageJ plugin). Then single stress fibers were manually picked up (the width of SF is set to 1µm) in ImageJ. The intensity of photoconverted actin (RFP channel) was normalized per intensity of total actin (GFP channel). The intensity changes were analyzed by a program in MATLAB written by Dr. ONG Hui Ting (Mechanobiology Institute, Singapore). The program dissected an isolated single SF into small squares (1µm× 1µm) and quantified the normalized intensity changes over time in those squares. The plateau maximal fluorescence intensity level and the time to reach half-maximal fluorescence intensity were calculated for each square.

### 4.4. Cyclic cell-stretching assays

The 3D-printed biaxial stretching device shown in Fig. 4A was developed by Dr. Li Qingsen (IFOM, Italy) (Li, 2018). A stretchable silicone membrane (thickness = 125µm) was coated with fibronectin. The stretching device was controlled by an ARDUINO UNO REV3 chip (ARDUINO, A000066). The software to input stretching programs into the chip was ARDUINO ver.1.8.9. The cells were seeded on the membrane the day before the experiment and the drugs were added 5min before stretching started. The membrane was cyclically stretched for 10% increase in diameter in 1Hz for 10min. Details of the time course of experiment with photoconverted actin incorporation upon stretching is shown in Figure 4B.

### 4.5. Traction force microscopy

The traction force microscopy with embedded beads is performed as described previously [29]. Briefly, a soft polydimethylsiloxane CY 52-276A and CY 52-276B (Dow Corning, 0008602722) were mixed with the ratio 1:1 and the Sylgard 184 crosslinker was used to tune the stiffness of the gel for proper force measurement of cells (∼95 kPa). The mixture was spin-coated onto a clean coverslip to achieve the thickness of ∼7μm and cured for 1 h at 80 °C. The surface of the gel was silanized with (3-aminopropyl) triethoxysilane for 2 h, followed by incubation of 0.04μm diameter carboxylate-modified dark red (660/680 nm) beads (Thermo Fisher Scientific, 1871942) at 1 X 10^6^ beads/ml in a solution of 0.1 M sodium bicarbonate for 30 min. Before seeding the cells, the coverslips with beads were further incubated for 30 min with 10 μg/ml fibronectin also dissolved in 0.1 M sodium bicarbonate and washed with Dulbecco’s phosphate-buffered saline (Sigma-Aldrich). The traction forces were calculated from bead displacement field visualized by live cell imaging as described in Tseng et al [30] using the online ImageJ plugin (https://sites.google.com/site/qingzongtseng/tfm for plugin software details). The computation algorithm by Sabass et al [87] was used. The distribution of traction force magnitude was presented as a heat map (Fig. 1E). The mean magnitude of the traction force values was calculated for each cell.

### 4.6. Fabrication of My-134 micropillars

PDMS micropillars were fabricated as described previously [31] to form PDMS mold for micropillar array. After silanizing the surface of the PDMS pillars with Trichloro (1H,1H,2H,2H-perfluorooctyl) silane (Sigma, 448931) overnight, new PDMS (DOWSIL 184 silicone elastomer, Dow Corning, MI, USA) was directly cast onto the surface of the micropillar to make a PDMS mold with holes. After degassing for 15 minutes, the mold was cured at 80 degree for 2 hours. The PDMS mold was peeled off from the PDMS pillars, cut 1cm square and placed on plastic dishes face up following a silanization of their surface with Trichloro (1H,1H,2H,2H-perfluorooctyl) silane overnight.

To fabricate the array of micropillar, whose refraction index is similar to that of the growth medium, a small drop of My-134 polymer (My Polymers Ltd., Israel) was put on the center of coverslip coated with 3-(Trimethoxyilyl)propyl methacrylate(sigma, 440159) and then, the silanized PDMS mold covered the drop face down onto the coverslip with thin layer of My-134 polymer for 15-30 min. After degassing for 5-15 min to get rid of air bubbles inside the polymer, the assembly was placed in a cell culture dish, covered with fresh milli-Q water and cured under short wavelength UV radiation (UVO Cleaner 342A-220, Jelight Company Inc, USA) for 6 min. Then, the PDMS mold was carefully peeled off from the coverslip.

Top of My-134 pillars were coated with fluorescence-labeled fibronectin as described previously (Doss et al., 2020). Briefly, human plasma fibronectin (Roche) was conjugated with Atto-647N using a protein labeling kit (cat # 76508 Sigma-Aldrich). PDMS stamps were incubated with solution containing 50μg/ml fibronectin and 1μg/ml of the conjugated fibronectin in Dulbecco’s phosphate-buffered saline (Sigma-Aldrich) at room temperature for 90 minutes. After washing with Milli-Q water and air-drying the surface, the PDMS stamp was put onto the top of My-134 pillars freshly exposed to UV-Ozone (UV Ozone ProCleaner Plus, BioForce Nanosciences). After 5 minutes of contact, the stamp was removed. Before cell plating, the My-134 pillars were incubated with 0.2% Pluronic F-127 (Sigma) for 1 hour for blocking, followed by washing three times with Dulbecco’s phosphate-buffered saline. The pillars in the array were arranged in a triangular lattice with 4 μm center-center distance and the dimensions of pillars were d=2.1μm with h=6μm (k = 50 nN/μm). The traction forces by fluorescent-labeled My-134 pillars were calculated using a custom-build MATLAB program (version 2019a, MathWorks) as described previously (Doss et al., 2020).

### 4.7. Actin and myosin flow measurement

REF52 cells transfected with vectors encoding photoconvertible PA-GFP-actin and tdTomato-MLC were plated overnight on fibronectin-coated 35 mm glass bottom dishes (Iwaki, 3930-035). Lines perpendicular to stress fibers were illuminated as the region of interest using MetaMorph software for 3-5 seconds by 150 cycles of illumination with 0.06mW 405nm laser for photoactivation, followed by image acquisition using W1 confocal unit equipped with Live-SR (SIM) system at 30 seconds interval for 15-30 minutes. This microscopy method permitted resolving of individual myosin filaments. For drug treatment, cells were pre-treated with 100 μM pAB or 25 μM SMIFH2 for 5 minutes before photoactivation.

To measure the velocity of movement of actin illuminated spots and myosin filaments in stress fibers, kymographs of the images of individual stress fibers were analyzed. Avarage speeds of the movement of actin spots or myosin filaments were calculated from the kymograph slope as shown in Figure 6B.

### 4.8. Statistical analyses

The methods of statistical analysis, the sample sizes (n) and p values are specified in the results sections or figure legends. Prism version 6 (GraphPad Software) was used to plot, analyze and represent the data. The quantitative data were presented in the figures as bar graphs or scatter dot plots showing mean ± SD.

## Acknowledgements

We are grateful to Ms. Hui Ting Ong (MBI, Singapore) for developing the software for data analysis, to Mr. Tang Hou Pang (MBI, Singapore) for technical support in My-134 micropillar fabrication method, to Drs. Andrea Ravasio and Jean-Francois Rupprecht (MBI, Singapore) for technical help with traction force microscopy and its analysis, to Dr. Bryant L. Doss (MBI, Singapore) for technical help with micropillar experiments and their analysis, and to Dr. Andrew Wong (MBI, Singapore) for kind help with editing of the manuscript. A.D.B. and V.V. acknowledge the support from the Singapore Ministry of Education Academic Research Fund Tier 2 (MOE Grant No: MOE2018-T2-2-138), the National Research Foundation, Prime Minister’s Office, Singapore, and the Ministry of Education under the Research Centres of Excellence programme through the Mechanobiology Institute, Singapore (ref no. R-714-006-006-271), and Singapore Ministry of Education Academic Research Fund Tier 3 MOE grant no. MOE2016-T3-1-002.

---

**Supplementary Figure1:**
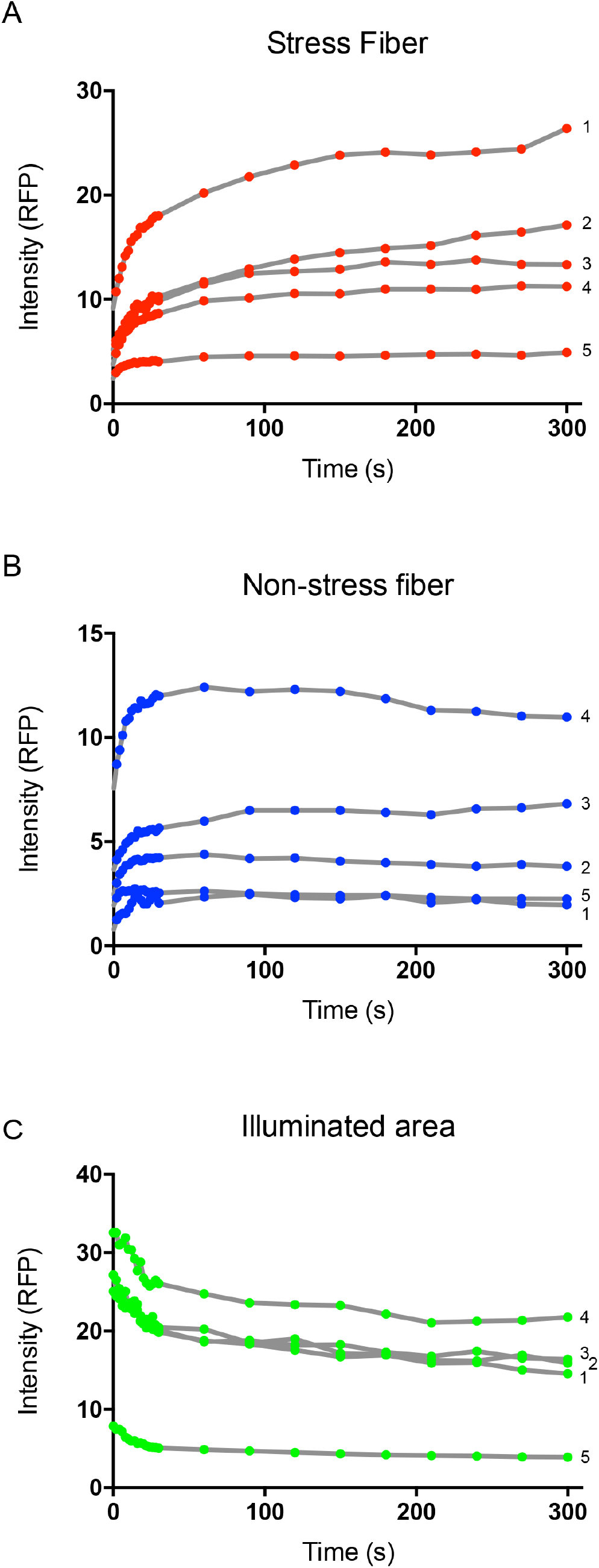
Graphs representing time course of fluorescence intensity dynamics of photoconverted actin (RFP channel) in the stress fiber region (A), the non-stress fiber area (B) and central illuminated area (C) of the cells shown in Figure 1C. See details in the legend to the Figure 1. Note that intensity of photoconverted actin fluorescence decreased in central illuminated area and increased in stress fiber and non-stress fiber regions.

**Supplementary Figure2:**
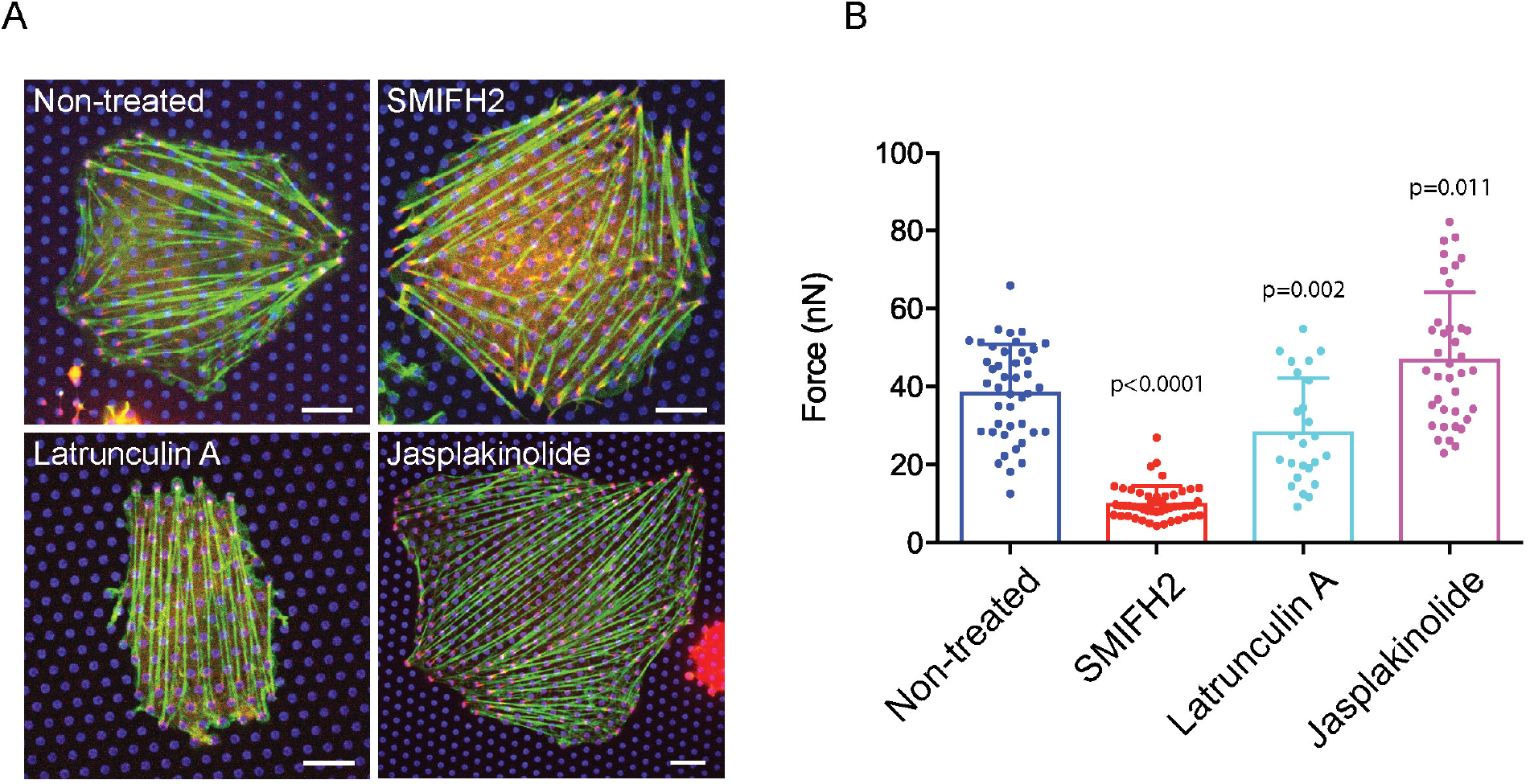
Traction forces generated by individual stress fibers as determined by micropillar deflection. (A) REF52 cells plated on micropillar array show stress fibers (green) and focal adhesions (red) labelled with GFP-β-actin and mApple-paxillin, respectively. Stress fibers are connected to micropillar tips labelled by Atto647N-fibronectin (blue). Non-treated cell (top, left) as well as cells treated with 30 μM SMIFH2 (top, right), 100 nM latrunculinA (bottom, left) or 100 nM jasplakinolide (bottom, right) are shown. Scale bar, 10μm. (B) The traction forces exerted by stress fibers on individual pillars in non-treated cells and cells treated with indicated drugs (25 µM SMIFH2 (n=43, N_cell_=9), 100 nM latrunculin A (n=25, N_cell_=4), and 100 nM jasplakinolide (n=38, N_cell_=5) for 30 min). Each dot represents the force value calculated from deflection of individual pillar. N: number of cells analyzed for each treatment, n: number of pillars measured. The bars represent the mean value over all pillars, the error bars show ± SD. The p-values characterizing the difference between mean values for non-treated and treated cells were calculated using an unpaired two-tailed student t-test.

**Supplementary Figure 3:**
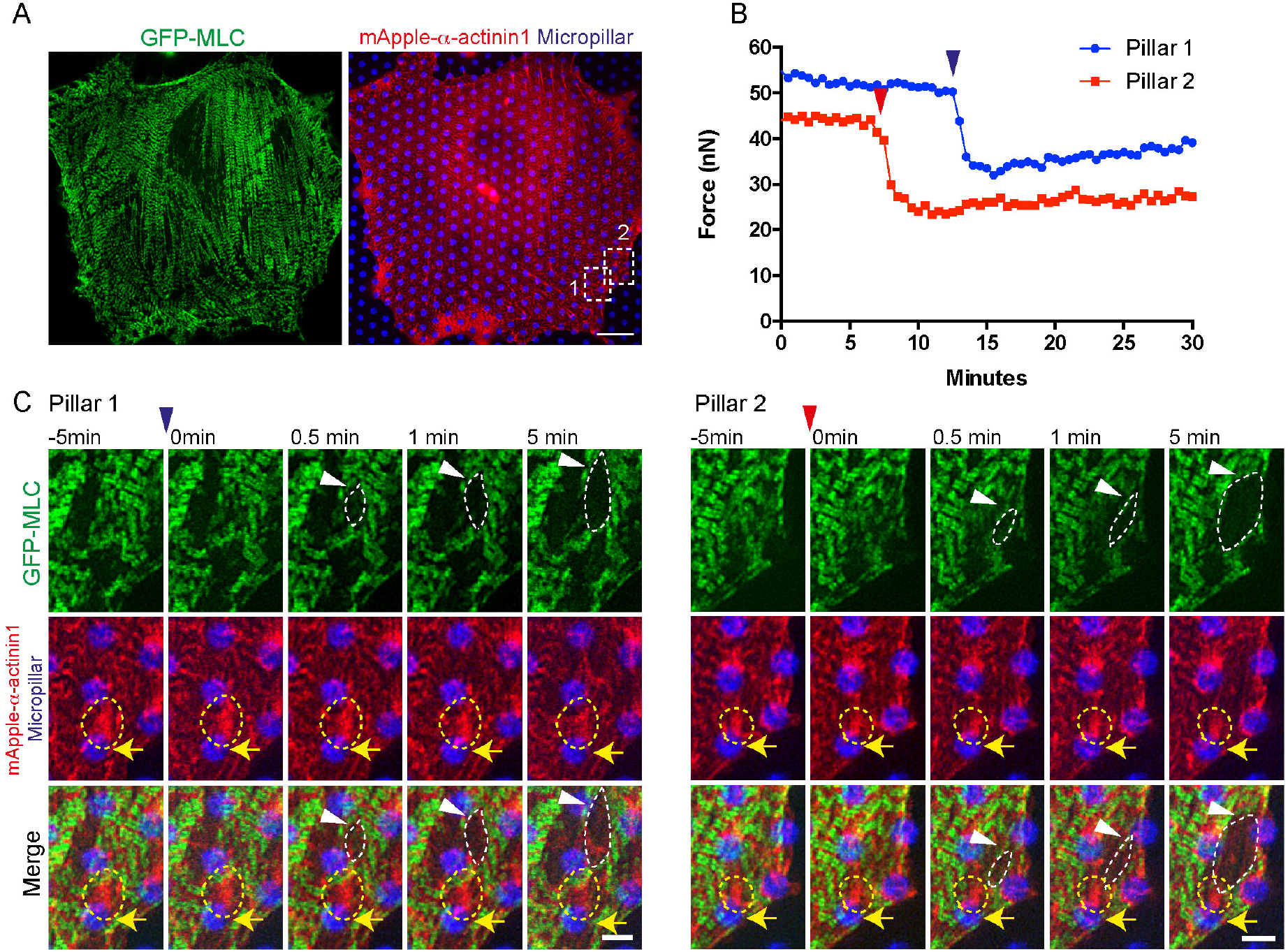
Latrunculin A induces local dropping of traction forces generated by individual stress fibers in the regions where “holes” in the α-actinin1/myosin II contractile network were detected. (A) REF52 cell plated on micropillar array preserved essentially intact myosin II filaments organized into stacks after treatment with 100 nM latrunculin A. Myosin II filaments (left image) were labelled with GFP-MLC (green), α-actinin1 striated distribution is shown in the right image of mApple-α-actinin1 (red), micropillar tips are labelled by Atto647N-fibronectin (blue). Scale bar, 10µm. (B) Two examples of measurements of traction forces in individual stress fibers of cell treated with latrunculin A. The forces calculated from the deflection of pillars in box1 (Pillar1, blue curve) and box2 (Pillar2, red curve) are shown. Blue and red arrowheads represent the time point of dropping traction forces, respectively. (C) Time course of the distribution of myosin II filaments and α-actinin1 in the boxes 1 (left panel) and 2 (right panel) at the high magnification. Upper row - myosin II filaments (green), middle row - a-actinin1 (red) and pillars (blue), and lower row - merged images. Latrunculin A treatment induces formation of oval ‘holes’ in the stacks of myosin filaments (marked by white dotted lines). Yellow arrowheads indicate the position of closest micropillar. Note that the drop of the traction forces shown in graph B correlates with appearance of these holes. The intensity of α-actinin1 overlapping with micropillars (yellow dotted circles) is decreased following formation of holes in myosin II distribution and drop of the traction forces, suggesting that disassembly of focal adhesions on micropillars follows the force dropping. Scale bar, 2µm.

## Movie legends

**Movie 1**

Incorporation of photoconverted actin into stress fibers and focal adhesions in REF52 cell. Movies of unconverted (GFP channel, left) and photoconverted mEos3.2-β-actin (RFP channel, right) are shown. The image acquisition starts just after 405 nm laser illumination of the circular area in the center of the cell and proceeded every 2-seconds over the first minute, and then every 30-seconds over a period of 5 minutes. Display rate is 10 frames/sec. The movie corresponds to the time-lapse series shown in Figure 1A. Scale bar, 20 μm.

**Movie 2**

The flow of myosin II filaments and photoconverted spots on actin filaments in stress fibers of REF52 cell. Myosin II filaments and photoconverted actin spots on stress fibers were visualized using tdTomato-MLC (green) and PA-GFP-β-actin (red), respectively. Myosin filaments are shown in the left frame, the photoconverted actin spots in the central frame and the merged images in the right frame. The frames were recorded immediately following actin photoconversion every 30-seconds over a period of 30 minutes using structured illumination microscopy (SIM). Display rate is 10 frames/sec. The movie corresponds to the kymographs shown in Figure 1D. Scale bar, 10 μm.

**Movie 3**

The dynamics of myosin II filaments and α-actinin1 in REF52 cell on micropillar array. SIM images of myosin light chain (left, GFP-MLC), α-actinin1 (middle, mApple-α-actinin 1), and merged (right, MLC in green; α-actinin1 in red; fluorescent fibronectin on pillars in blue) are shown. The frames were recorded every 30-seconds over a period of 30 minutes. Display rate is 10 frames/sec. The movie corresponds to the images shown in Figure 1F. Scale bar, 10 μm.

**Movie 4**

The dynamics of traction forces, calculated from pillars displacements, and myosin II filaments in REF52 cell. Myosin II filaments (GFP-MLC, green), pillars (fluorescent fibronectin, magenta) and traction force vectors (yellow) are shown. The frames were recorded every 30-seconds over a period of 30 minutes using SIM. The movie corresponds to the image shown in Figure 1G.

**Movie 5**

REF52 cell treated with 100 nM latrunculin A on micropillar array. Myosin II filaments labelled with GFP-MLC (green) are shown in the left frame, α-actinin 1 labelled with mApple-α-actinin1 (red) and micropillar tips labelled by Atto 647N-fibronectin (blue) are shown in the right frame. The frames were recorded immediately following the latrunculin A addition, every 30-seconds over a period of 30 minutes using SIM. Display rate is 10 frames/sec. The movie corresponds to the time-lapse series shown in Supplementary Figure 3A. Scale bar, 10 µm.

**Movie 6**

The flow of myosin II filaments (green, tdTomato-MLC) and actin filaments (red, photoconverted PA-GFP-β-actin) in individual stress fibers of non-treated (top), and treated with 100 μM pAB (middle) and 25 μM SMIFH2 (bottom) cells. Focal adhesions are located on the left end of the stress fibers. The frames were recorded every 90-seconds over a period of 30 minutes for non-treated cell and every 45-seconds over a period of 15 minutes for both pAB and SMIFH2 treated cells. Display rate is 10 frames/sec. The movies correspond to the kymographs shown in Figure 6A. Scale bar, 2 µm.

**Movie 7**

The flow of myosin filaments (green, tdTomato-MLC) in REF52 cell treated with 100 μM pAB on micropillars (blue, Atto 647N-fibronectin). The distribution of focal adhesions (red, GFP-vinculin) is shown in the first and the last frame. The frames were recorded immediately following the drug addition every 30-seconds over a period of 30 minutes. Display rate is 10 frames/sec. The movie corresponds to the kymograph shown in Figure 6C. Scale bar, 10 µm.

**Movie 8**

The flow of myosin filaments (green, tdTomato-MLC) in REF52 cells treated with 25 μM SMIFH2 on micropillars (blue, Atto 647N-fibronectin). The focal adhesions (red, GFP-vinculin) are shown in the first and the last frame. The frames were recorded immediately following the drug addition every 30-seconds over a period of 30 minutes. Display rate is 10 frames/sec. The movie corresponds to the kymograph shown in Figure 6D. Scale bar, 10 µm.

**Movie 9**

The flow of myosin filaments (green, tdTomato-MLC) in REF52 cells treated with the mixture of 100 μM pAB and 25 μM SMIFH2 on micropillars (blue, Atto 647N-fibronectin). The focal adhesions (red, GFP-vinculin) are shown in the first and the last frame. The frames were recorded immediately following the drug addition every 30-seconds over a period of 30 minutes. Display rate is 10 frames/sec. The movie corresponds to the kymograph shown in Figure 6E. Scale bar, 10 µm.

